## Supplementary figures and images for "Neuron-specific RNA-sequencing reveals different responses in peripheral neurons after nerve injury"

### Figure 1 Supplement 1

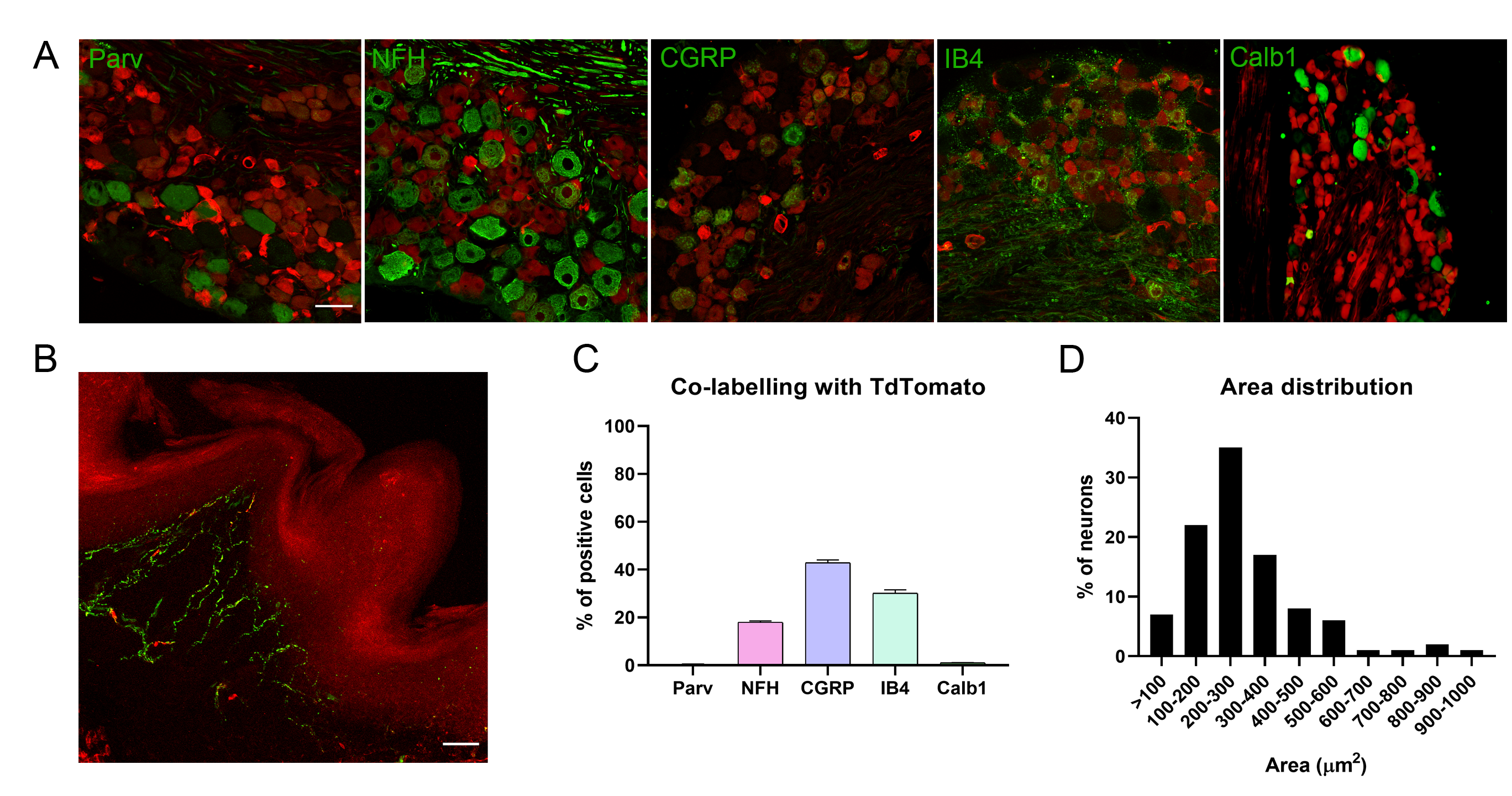

### Figure 1 supplement 2

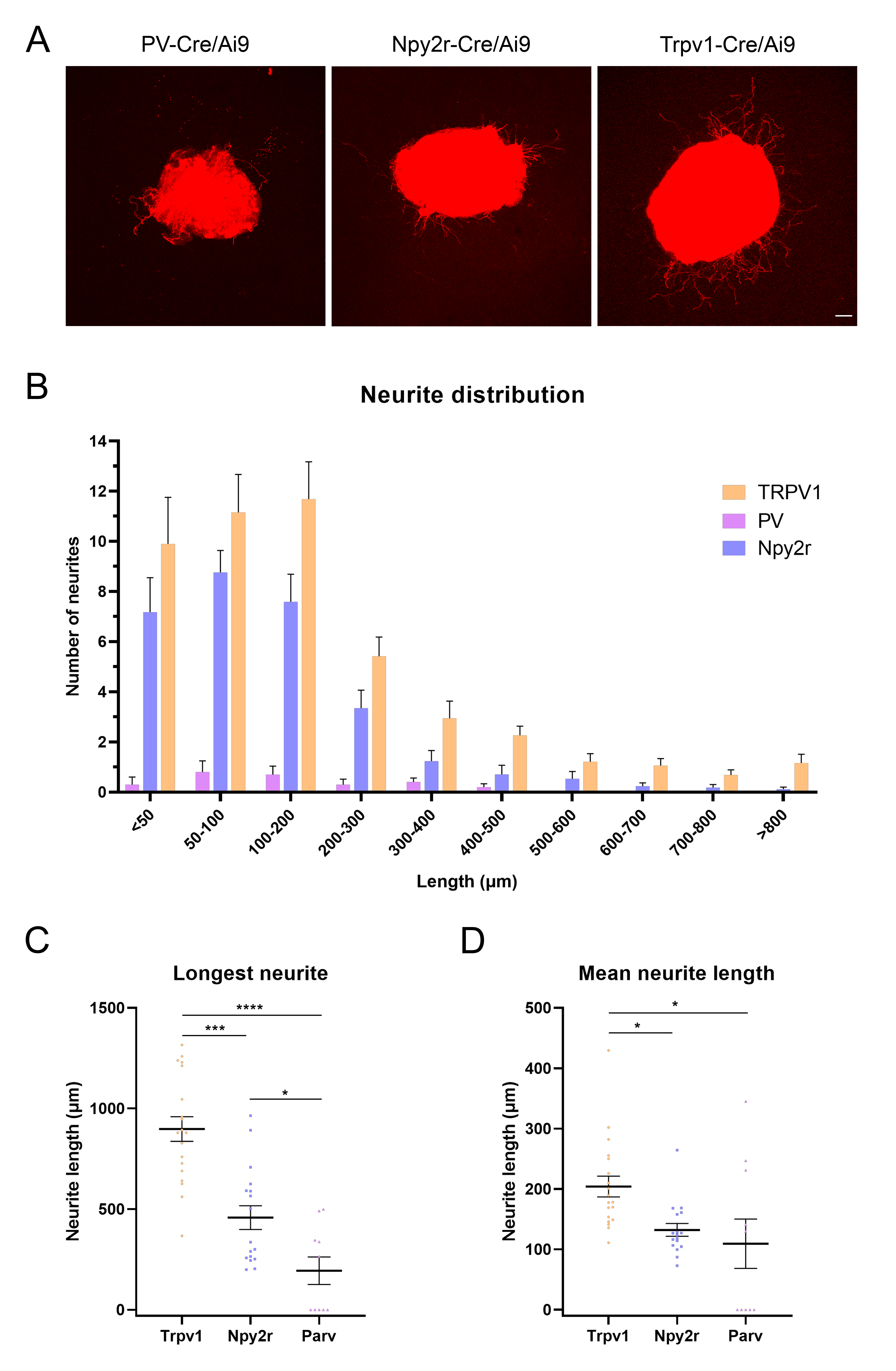

### Figure 2 supplement 1

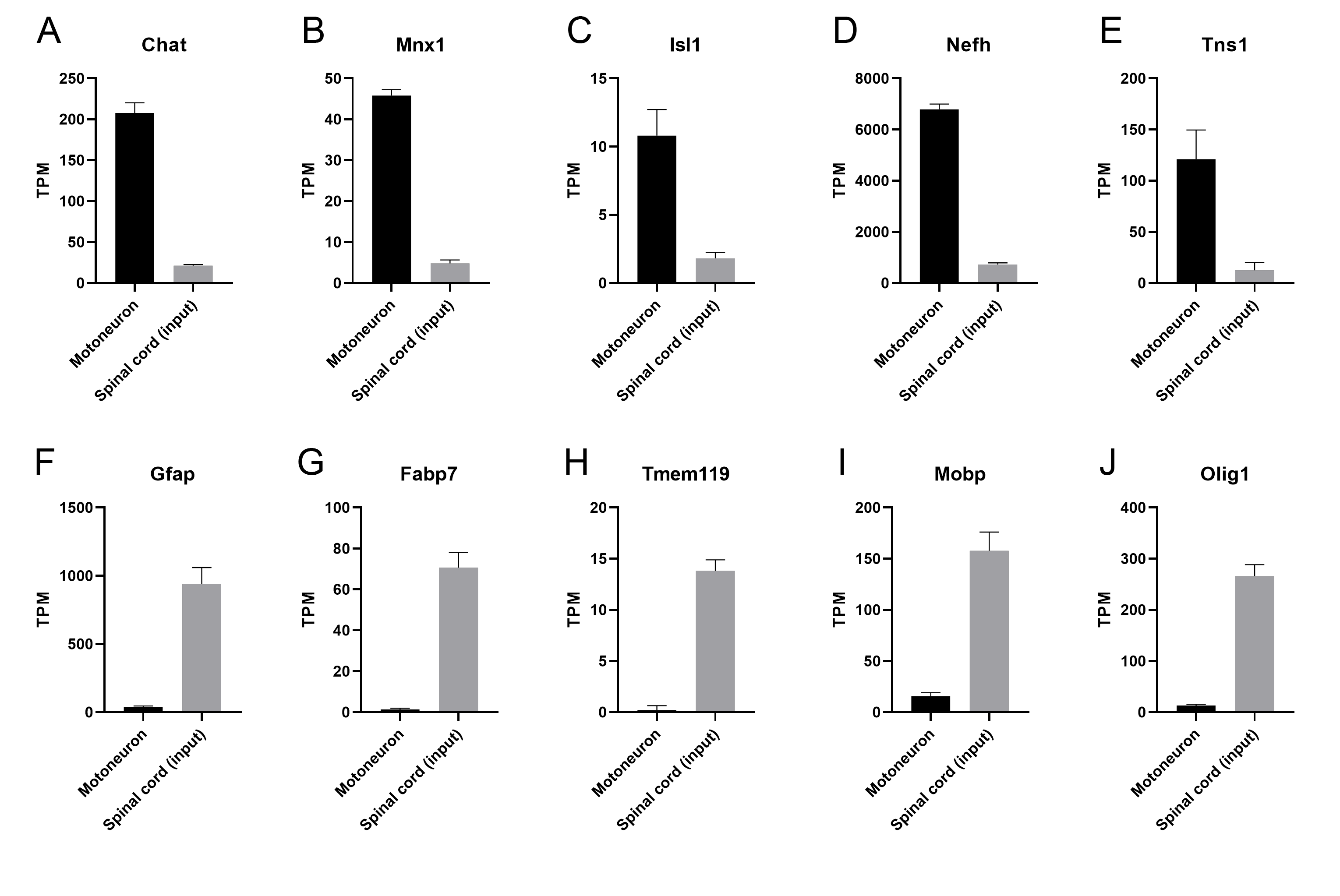

### Figure 2 Supplement 2

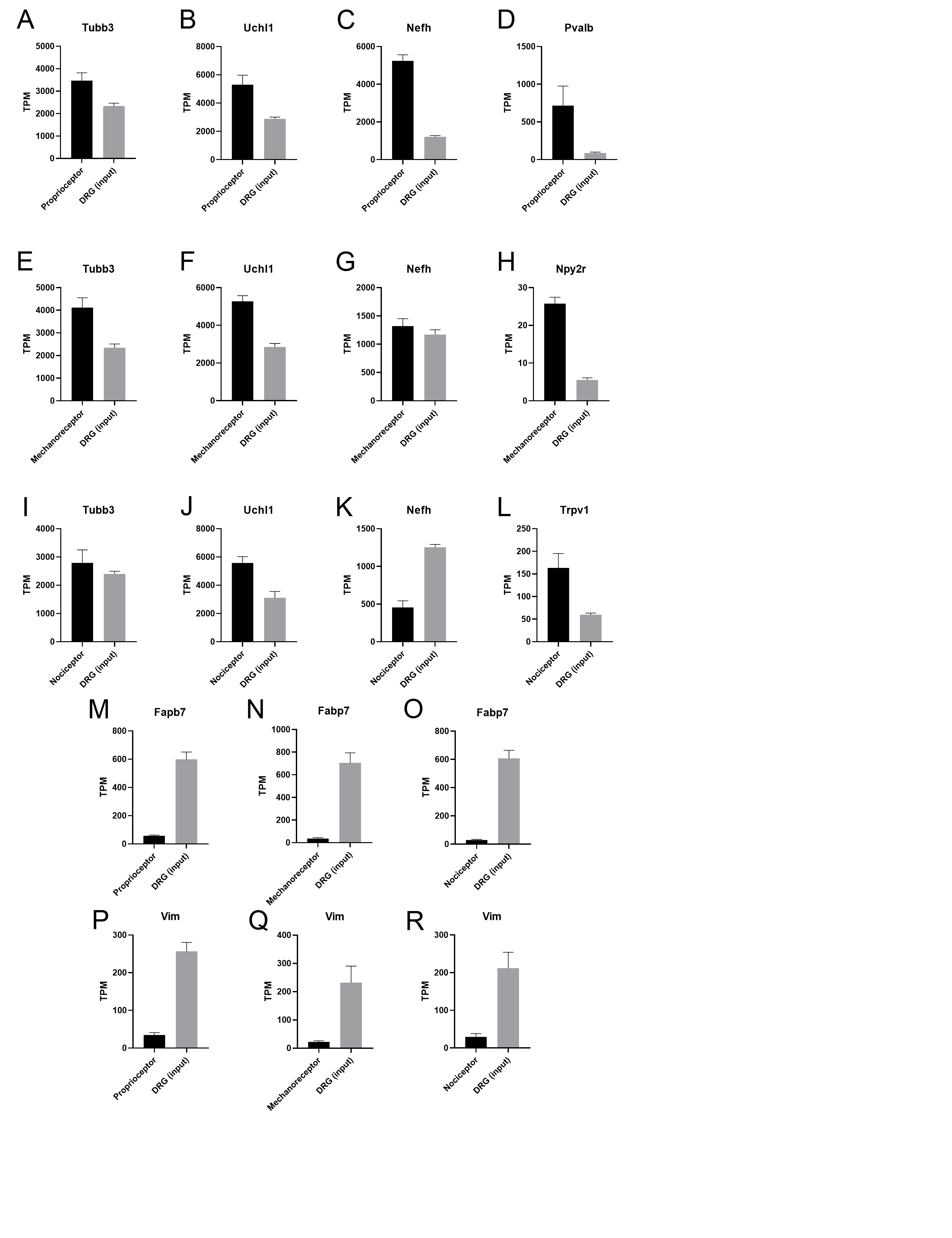

### Figure 2 supplement 3

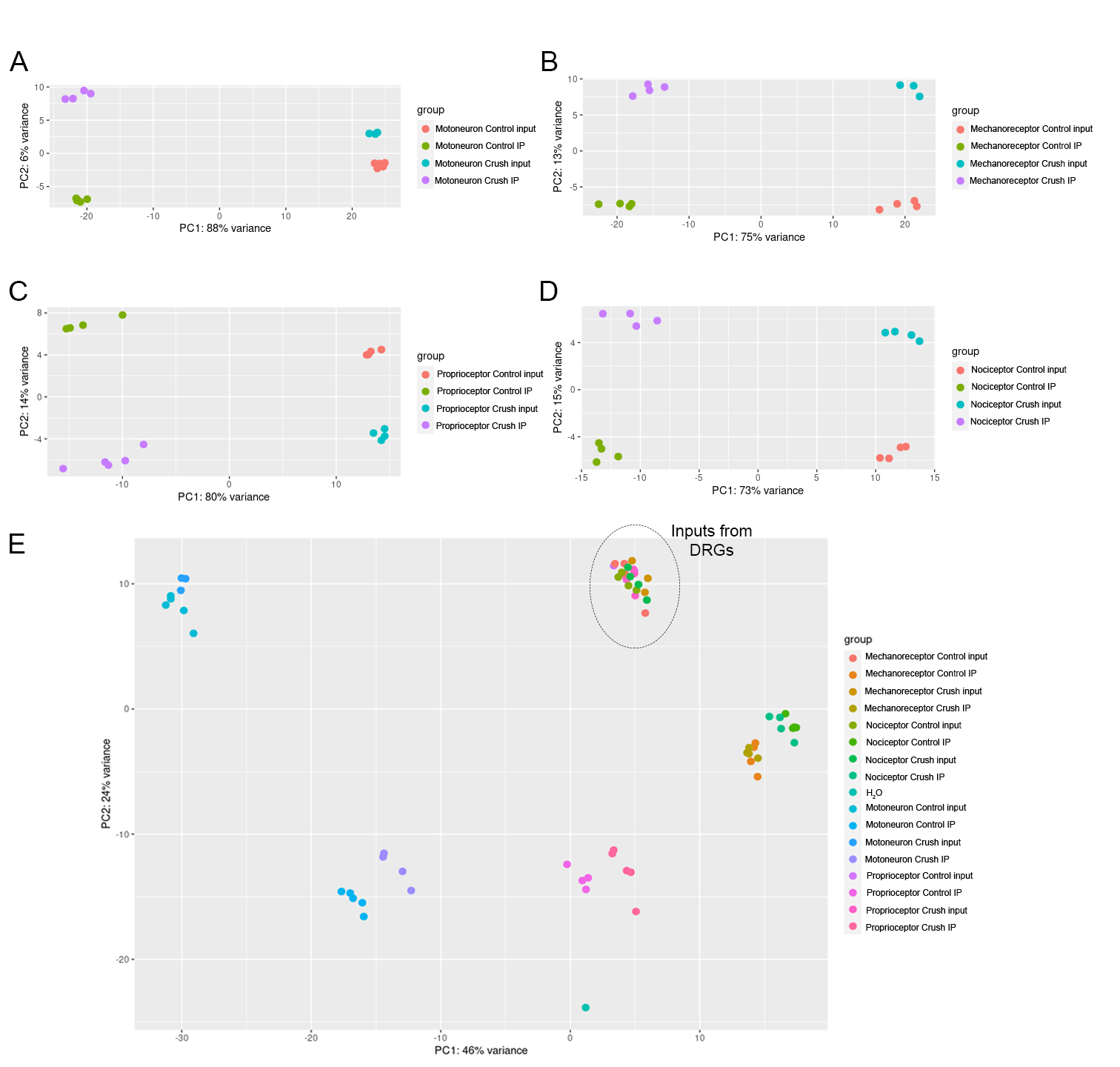

### Figure 2 supplement 4

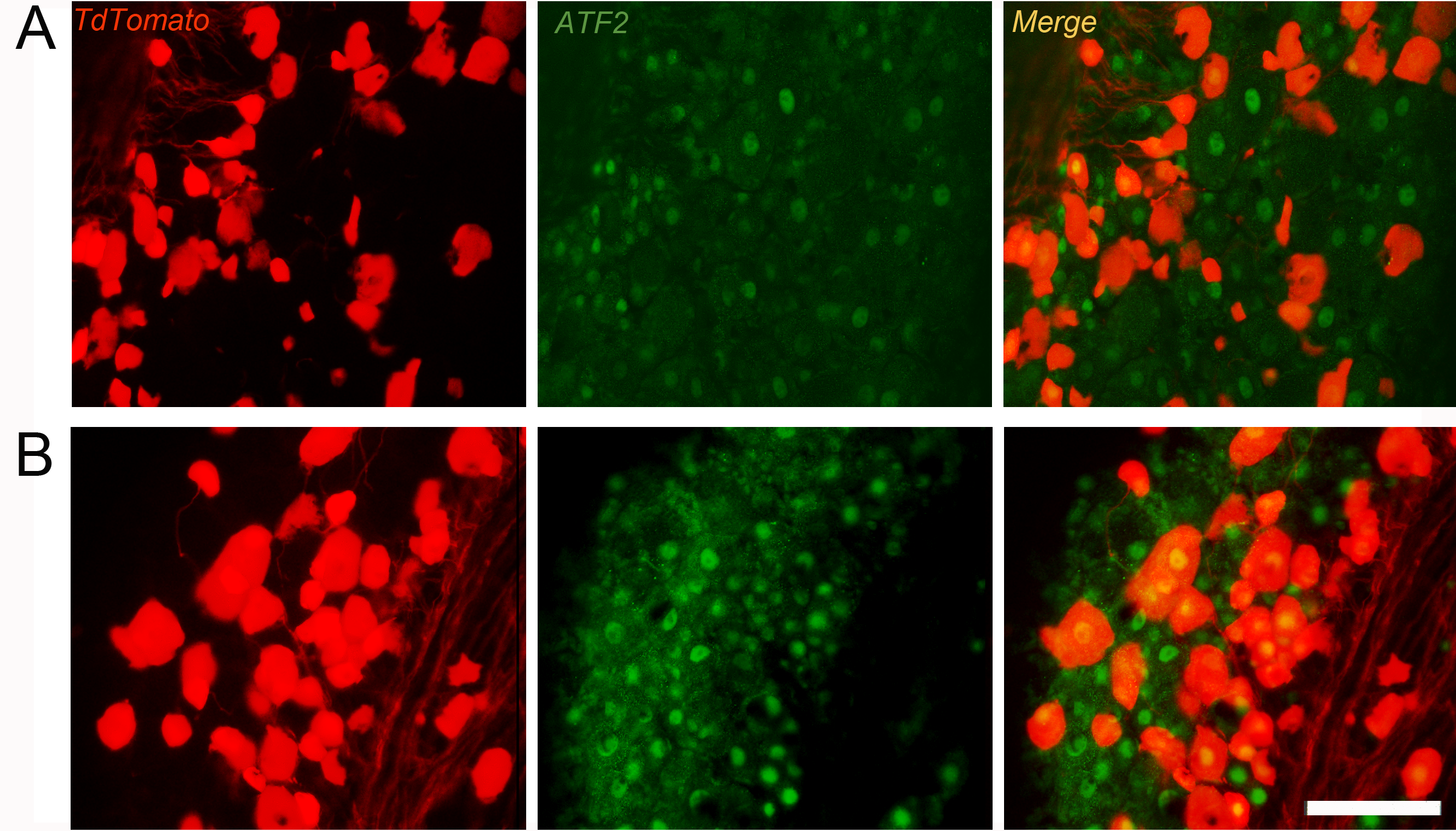

### Figure 3 supplement 1

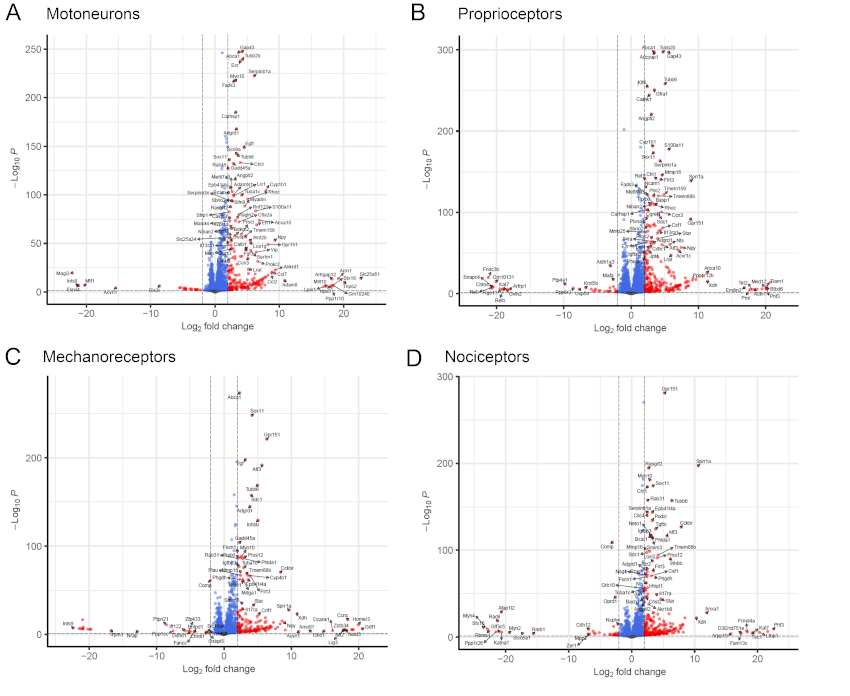

### Figure 3 supplement 2

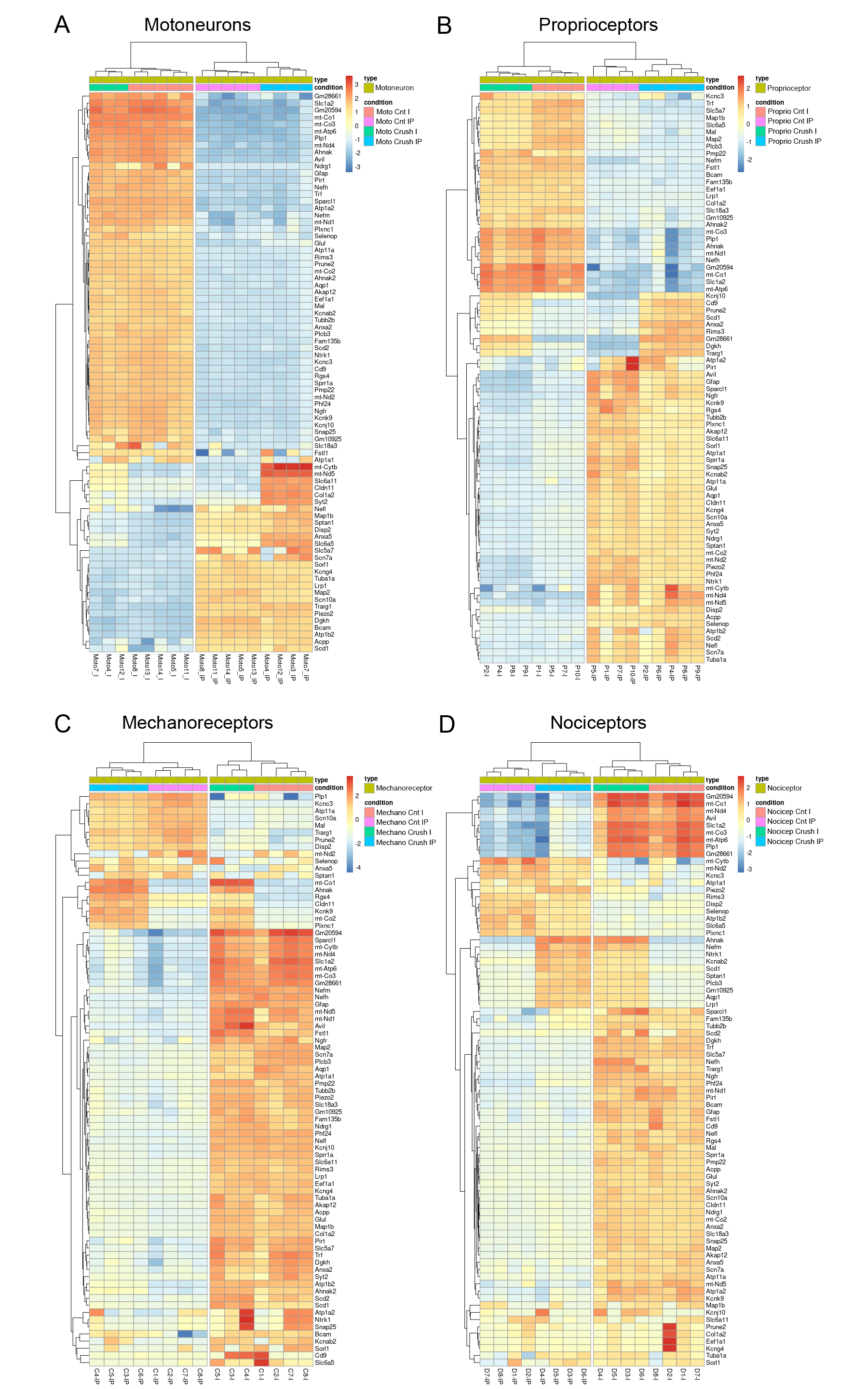

### Figure 3 supplement 3

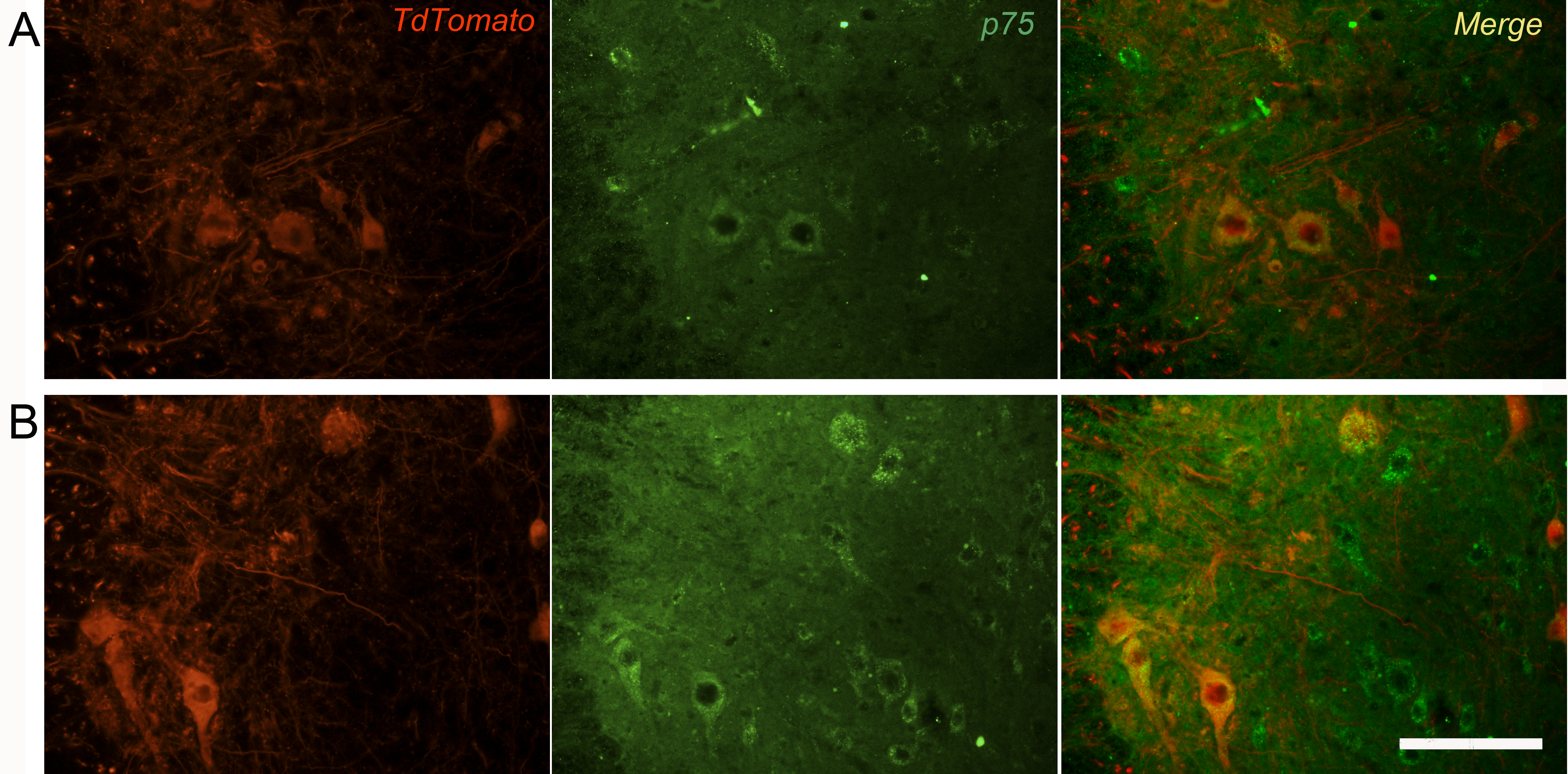

### Figure 5 supplement 1

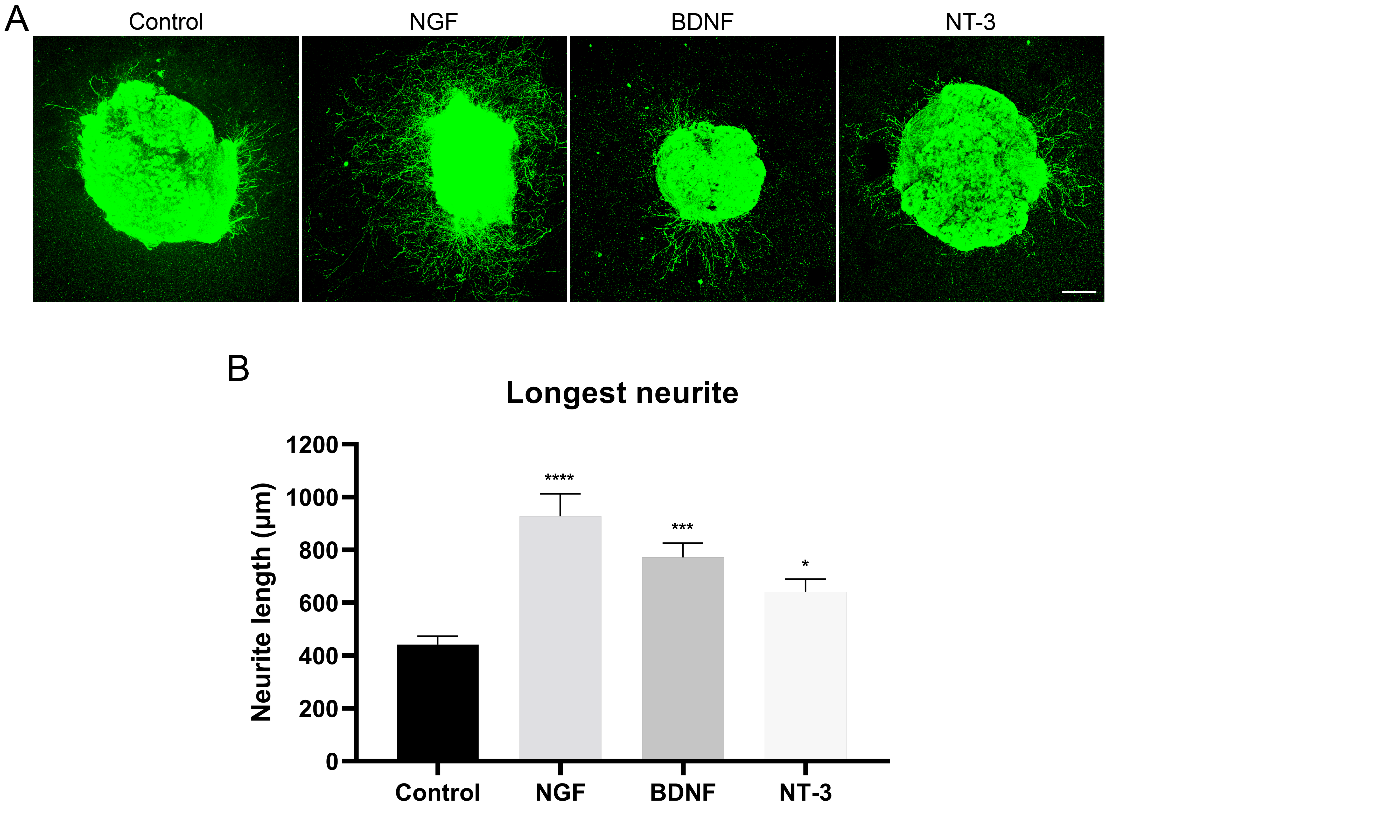

### Figure 6 supplement 1

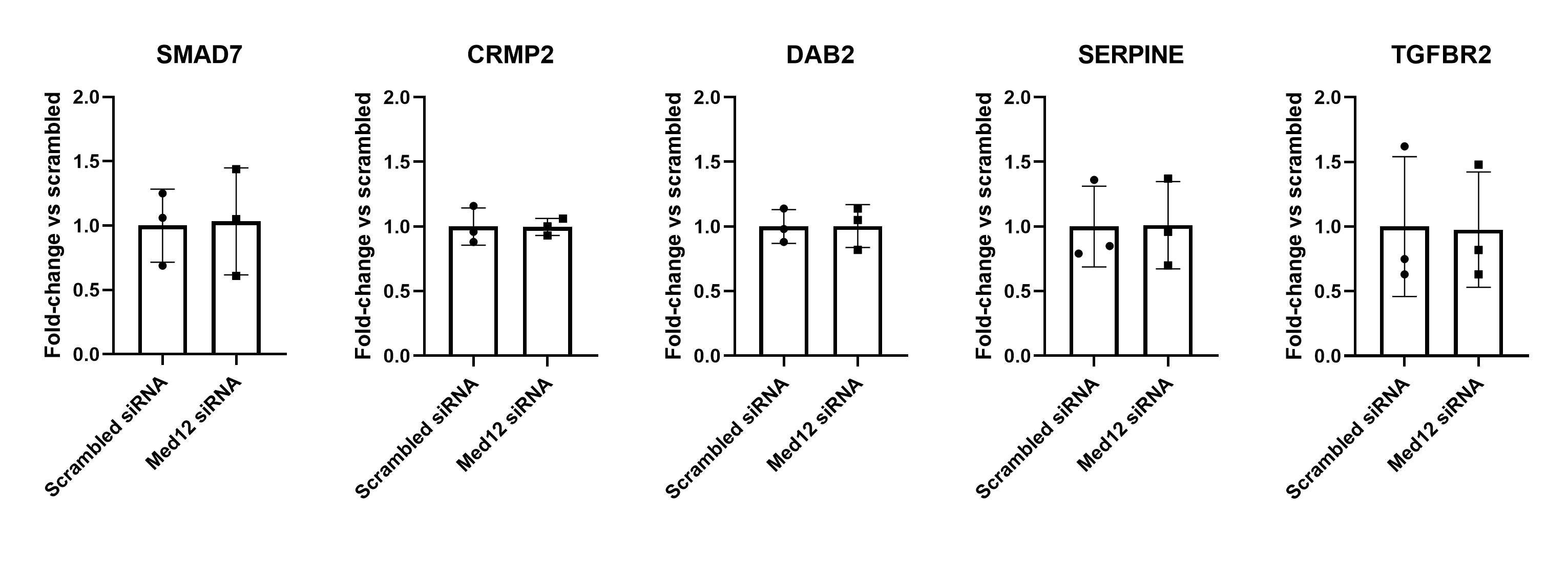
